# Eukaryotic ribosome recognizes 3′ context and stimulates stop codon readthrough

**DOI:** 10.1101/2021.04.18.440353

**Authors:** Elizaveta Sokolova, Tatiana Egorova, Alexey Shuvalov, Elena Alkalaeva

## Abstract

It is known that the nucleotide context surrounding stop codons significantly affects the efficiency of translation termination. In eukaryotes, various 3′ contexts have been described that are unfavourable for translation termination; however, the exact molecular mechanism that mediates their effect remains unknown. In this study, we used a reconstituted mammalian translation system to examine the efficiency of stop codons in different contexts, including several previously described weak 3′ stop codon contexts. Our results revealed that ribosomes can independently recognize certain contexts and ignore stop codons that are followed by these sequences. Moreover, the efficiency of translation termination at the weak 3′ contexts was almost equal to the one at the standard context. We propose that weak 3′ contexts interact with the 18S rRNA provoking a conformational change in the U-turn-like structure of the stop codon in the A site of ribosome. This change makes incorporation of the near-cognate tRNA more preferable than recognition of the stop codon by the release factors and increases readthrough.

## INTRODUCTION

The protein synthesis ends when a termination signal sequence of mRNA containing a stop codon (UAA, UAG or UGA) occupies the ribosome’s A-site, where protein release factors decode it. In eukaryotes, tRNA-mimicking factor eRF1 recognizes all three stop codons and promotes release of the synthesized peptide from the peptidyl-transferase center. It is stimulated by the GTP-ase factor eRF3, which resembles the elongation factor eEF1A (1–4). Remarkably, there are positional and conformational similarities between eRF1-eRF3 and elongation aa-tRNA-EF-Tu complexes (5, 6). Stop codon recognition is implemented by the conserved TASNIKS and YxCxxxF motifs of the N-domain of eRF1 (7, 8). The lysine of NIKS motif can be hydroxylated, which improves termination efficiency (9). Cryo-EM structures of mammalian ribosomal complexes containing stop codon in the A-site have shown that binding of eRF1 leads to changes of mRNA configuration so that the fourth nucleotide following three bases of stop codon is pulled into the A-site (10–12). This configuration differs from the shape of sense codons, recognized by tRNAs, and implicates complex three-dimensional interplay of the release factor and 18S rRNA. It also explains the strong impact of the identity of 3’ nucleotide downstream of stop codon on termination (13).

However, sometimes the amino acid is incorporated into the nascent polypeptide chain instead of proper translation termination. Such an event is a result of stop codon suppression, or readthrough, when the stop signal in the ribosomal A-site is interpreted as a sense codon and is recognized by near-cognate tRNA rather than by the release factor. The basal level of readthrough on naturally occurring stop codons commonly has a frequency of < 0.1% (14), although readthrough rates in some cases recently were shown to be higher than 10% (15, 16). The phylogenetic study of 12 Drosophila species revealed more than 280 cases of conserved stop codon readthrough. It was confirmed with ribosome profiling analysis, which indicated the readthrough as a relatively common event (17, 18). Subsequent researches identified readthrough in fungi (19), and numerous works continue to describe readthrough for a large number of transcripts in mammals (15, 16, 20–24). Besides, premature stop codons also sustain quite a sufficient percent of readthrough (<1%) (14), which is essential for therapeutics of the diseases, occurring as a result of nonsense mutations. All these data demonstrate the importance of readthrough events, which can be considered not only as an error during termination process but rather as an important regulatory mechanism.

It was shown that the nucleotide context, surrounding stop codons, significantly affects the level of readthrough in different groups of eukaryotes, pointing to possible context-driven control of protein synthesis, its localization, and function, which can regulate the organismal state (25–27). 5’ and 3’ nearest environs of stop codons can decrease and increase translation termination efficiency (28). Earlier, the most strong influence on translation termination was demonstrated for +4 nucleotide, immediately following stop codon (13, 29–31). The statistical analysis of more than 5000 genes in mammals (13) with further *in vitro* experiments showed termination efficiency change according to context, which allowed to formulate the rule: for all three stop codons +4 A=G>>U=C, with the frequency of termination signal descended in the row: UAA(A>G), UAG(G>A), UGA(G>A), and the least – UAG(C/U), UGA(C/U). Authors of another work (32) estimated the efficiency of stop codon suppression in lacZ reporter system in *S. cerevisiae* and came to a slightly different conclusion: termination efficiency decreases among UAA G>A>U>C, UGA G>U>A>C and UAG A>U>C>G. Few work investigating an influence of 3’ context allowed to propose that terminating signal includes four (33, 34), or six nucleotides (35, 36). There is a preference for purines to pyrimidines in eukaryotic genomes (37, 38). Among searched genes of different taxonomic representatives, *D. melanogaster, H. sapiens, C. elegans, A. thaliana, S. cerevisiae*, the most frequent nucleotide in +4 position is A or G (38). The more comprehensive analysis of genomes of diverse eukaryotic groups (vertebrates, invertebrates, mono- and dicotyledons, yeast, and protozoans) with the use of statistical approach showed, that depending on stop codon there are insignificant differences in prevalence of either nucleotide in +4 position between single groups, but with the apparent preference of purines (37). In eukaryotes as a whole, the most widespread are UAA(A/G), and UGA(A/G) stop signals. For example, in *S. cerevisiae* and *D. melanogaster* A/G in +4 is preferable, and their genes with high expression levels have UAAG stop sequences (33). Nucleotide distribution up to +9 in *S. cerevisiae* and most likely in all eukaryotes is not random. The importance of at least six nucleotides following stop codon for readthrough was shown in *S. cerevisiae*. The *in vivo* analysis (36) denotes that positions +4, +5, +6, +8, and +9 are the key, at that +7 position did not have any effect. 3’ context CA(A/G)N(C/G/U)A discovered in this search just slightly differed from the corresponding consensus 3’ sequence from tobacco mosaic virus (TMV). The most effective suppression motif CAA UUA, revealed in this work, entirely conforms to the 3’ context of stop codon UAG of TMV. Another investigation in yeast proved the dependence of readthrough on +4, +5, +6, +8 (or +9) positions of 3’ context (39). Cridge and co-workers (40) affirmed the high impact of +4 and +8 nucleotides downstream of each stop codon on the readthrough in mammalian cells. They showed that +5 and +6 positions determined the increase or decrease of readthrough depending on the stop codon and +4 nucleotide identity. It points to the combined effect of neighboring nucleotides, and the uniqueness of +7 and +9 nucleotides was not significantly influential.

There is evidence of factors, which can influence the readthrough levels in cooperation with the stop codon context. It was shown that translation initiation factor eIF3 increases readthrough on weak termination contexts, possibly promoting the incorporation of near-cognate tRNAs (41). The posttranslational hydroxylation of prolyl in ribosomal protein Rps23 of the 40S subunit can also modulate termination accuracy in a context-dependent manner (42). However, the exact mechanism of the context influence on translation termination remains unknown yet.

According to our previous work (43), there was no apparent connection between nucleotide frequencies in the 3’ stop codon contexts and their effect on peptide release efficiency. To reveal the mechanism of the context influence on termination, we tested known weak stop codon 3′ contexts in reconstituted translation termination.

## MATERIALS AND METHODS

### Construction of model mRNAs

MVHL-stop mRNAs containing one of the three stop codons (UAA, UAG, or UGA) were described previously (3, 44). To obtain mRNAs with the desired 3′ stop codon contexts and with the second UAA stop codon following the context of interest, site-directed mutagenesis was carried out. Primers for each construction are listed in the Supplementary Data (Table S1, S2). For run-off transcriptions, all plasmids were linearized with XhoI. mRNAs for pre-termination complex (preTCs) reconstruction were transcribed by T7 RNA polymerase using run-off transcription.

### Ribosomal subunits and translation factors

40S and 60S ribosomal subunits, as well as eukaryotic translation factors eIF2, eIF3, eEF1, and eEF2, were purified from a HeLa cell lysate, as described previously (3). The eukaryotic translation factors eIF1, eIF1A, eIF4A, eIF4B, ΔeIF4G, ΔeIF5B, eIF5, eRF1, mutant eRF1(AGQ), and eRF3c, lacking the N-terminal 138 amino acid residues, were produced as recombinant proteins in *Escherichia coli* strain BL21 with subsequent protein purification using Ni-NTA agarose and ion-exchange chromatography (3). Human full-sized eukaryotic release factor eRF3a (GSPT1) was kindly provided by Dr. Christiane Schaffitzel and was expressed in insect cell line Sf21 with baculovirus EMBacY from a MultiBac expression system (45).

### PreTCs assembly in vitro

Initiation complexes were assembled at 4°C and contained 2 pmol mRNA, 6 pmol Met-tRNAi^Met^, 4.5 pmol each of 40S and 60S ribosomal subunits, 7.5 pmol each of eIF2, eIF3, eIF4A, ΔeIF4G, eIF4B, eIF1, eIF1A, eIF5, and ΔeIF5B, supplemented with buffer composed of 20 mM Tris acetate, pH 7.5, 100 mM KAc, 2.5 mM MgCl_2_, 2 mM DTT, 0.3 U/µL RNAse inhibitor, 1 mM ATP, 0.25 mM spermidine, and 0.2 mM GTP to reach a final volume of 30 µL. The reaction mixture was kept at 37°C for 15 min to allow ribosomal-mRNA complex formation, and a 10 µL aliquot was subsequently subjected to a toe-print assay. Peptide elongation was performed on the remaining 20 µL by the addition of 3 pmol total tRNA (acylated with all or individual amino acids), 8 pmol eEF1, and 2 pmol eEF2 to the initiation complex and was incubated for 15 min at 37°C. A 10 µL sample containing preTC was subsequently subjected to a toe-print assay. The last 10 µL sample containing preTC was supplemented with 5 pmol eRF1 and 5 pmol eRF3a/c. The reaction mixture was incubated at 37°C for 15 min and subsequently subjected to a toe-print assay.

### Purification of preTCs and translation termination assays

A preparative amount of preTC was assembled *in vitro*, as described previously (46), and used in a conformational rearrangement analysis (47, 48). Briefly, 37 pmol of mRNA was incubated for 30 min at 37°C in buffer A (20 mM Tris acetate, pH 7.5, 100 mM KAc, 2.5 mM MgCl_2_, 2 mM DTT) supplemented with 400 U RNAse inhibitor, 1 mM ATP, 0.25 mM spermidine, 0.2 mM GTP, 75 µg total tRNA (acylated with all or individual amino acids), 75 pmol purified 40S and 60S ribosomal subunits, 125 pmol each eIF2, eIF3, eIF4A, ΔeIF4G, eIF4B, eIF1, eIF1A, eIF5, ΔeIF5B, 200 pmol eEF1, and 50 pmol eEF2. Next, the mixture was centrifuged in a Beckman SW55 rotor for 95 min at 4°C, and 50000 rpm in a 10–30% (w/w) linear sucrose density gradient (SDG) prepared in buffer A with 5 mM MgCl_2_. According to optical density, fractions corresponding to preTC complexes were combined, diluted 3-fold with buffer A containing 1.25 mM MgCl_2_ (final concentration 2.5 mM Mg^2+^), and analyzed via a toe-printing assay. For that, 10 µL aliquots containing 0.03 pmol preTCs were incubated at 37°C for 15 min with 1 µL buffer A or 0.625 pmol of eRF1(AGQ) and eRF3c/a with 0.2 mM GTP supplemented by 0.2 mM Mg^2+^. Samples were analyzed using a primer extension protocol. Toe-printing analyses were performed with AMV reverse transcriptase and a 5’
s-FAM labeled primer (5’-FAM-GCATTTGCAGAGGACAGG-3’) complementary to β-globin mRNA nucleotides 197–214. cDNA was separated by electrophoresis using standard GeneScan® conditions on an ABI Prism® Genetic Analyser 3100 (Applera).

The level of basal readthrough was calculated using rfu signals for ribosomal complexes in the formula 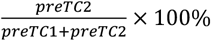. The percent of terminating ribosomes at the first stop codon was calculated using the formula 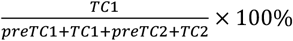.

## RESULTS

### Ribosome recognizes 3’ stop codon context independently of eRFs

The context from MVHL-stop mRNA, UGU CGU, was chosen as standard. This context we used in all previous studies in reconstituted mammalian translation system (3, 43, 44, 46, 49, 50). The frequencies of triplets from this sequence are close to that calculated for random triplet NNN 1/64 = 1.56% (43). Thus, this sequence can be legitimately used as a control one (Table 1).

**Table 1.** Nucleotide sequences of the selected 3′ contexts.

|  | Stop codon | 3' context |
| --- | --- | --- |
| Standard | UAA | UGU GUG |
|  | UAG |  |
|  | UGA |  |
| Weak 1 | UGA | CUA GUA |
| Weak 2 | UGA | CUA UAC |
| Weak 3 | UGA | CUU AAA |
| Dyst | UAG | GAU AAU |
| TMV | UAG | CAA UUA |

The consensus sequence UGA CUAG from a few mammalian genes was previously shown to be preferable for readthrough to occur (15). Therefore, we cloned two sequences, CUA GUA (Weak 1), and CUA UAC (Weak 2) and inserted them after the UGA stop codon (Figure 1A, Table 1). In the human genome, CUA GUA follows the UGA stop codon in the two genes *DDX58* and *D102*, while CUA UAC follows the UGA stop codon in the gene *CCNH*. preTCs were obtained as described above. We then performed toe-print analyses of the ribosomal complexes in order to assess preTC formation. Normally in the presence of standard and other tested contexts, preTCs are detected in the toe-print assay as a single peak at 15 nucleotides after the stop codon (Figure 1B). For both the weak contexts (CUA GUA and CUA UAC after the UGA stop codon), we observed an additional peak corresponding to a +3 nucleotide forward shift of the ribosomal complex from the UGA stop codon, indicating an additional translation elongation step (Figure 1B). We propose that the ribosome recognizes the 3′ end of the stop codon independently and is able to suppress stop codons using near-cognate tRNA in the absence of release factors. Notably, the +3 nucleotide shift of the ribosomal complexes occurred only in the presence of aminoacylated total calf tRNA, while addition of other aminoacylated tRNAs (MVHL tRNA transcripts or total yeast tRNA) to the translation elongation did not provoke a +3 shifted peak (Figure 2). It appears that only mammalian total tRNA contains suppressor or near-cognate tRNA suited to the ribosomal A site containing the UGA stop codon. It is well known that near-cognate tRNAs are able to suppress UGA stop codon in yeast (51, 52). We suppose that there is no contradiction with our data, as the ribosomes used in our experiments were mammalian. Possibly, yeast tRNAs don’t work as near-cognate in mammalian translation system. The reason why we observed that the ribosome performed only one translocation step might be depletion of tRNA mix. Indeed, the ratio of aminoacylated calf tRNAs is different. That’s why in further experiments we added the higher amount of tRNAs mix.

**Figure 1.**
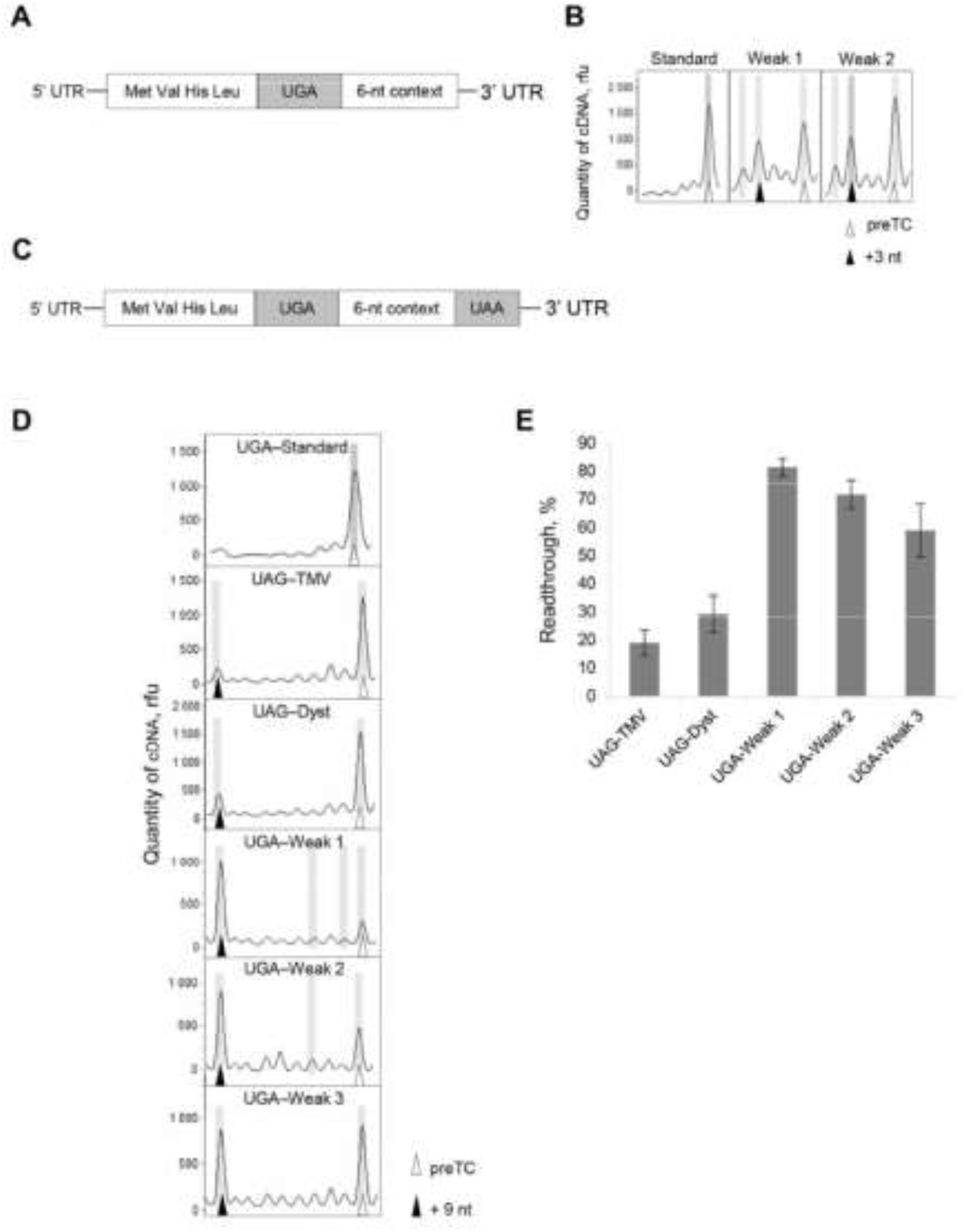
Ribosome detects a 3′ stop codon context independent of eRFs. (A) The scheme of the model mRNA carrying various stop codon contexts. (B) Toe-print analysis of the ribosomal complexes assembled with mRNAs containing various stop codon contexts. (C) The scheme of the dual-stop mRNA, used for quantification of readthrough levels. (D) Toe-print analysis of ribosomal complexes assembled at the dual-stop mRNAs containing various stop codon contexts. (E) Quantification of readthrough levels obtained with the dual-stop mRNAs. rfu, relative fluorescence unit; n, number of repeats.

**Figure 2.**
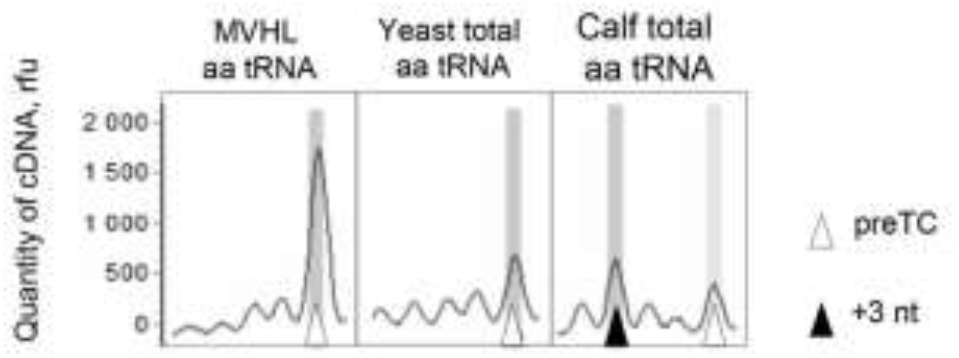
Translational readthrough in the presence of total aminoacylated tRNAs from different sources. Example of toe-print analysis of pretermination complexes assembled in the presence of calf and yeast total tRNA, or MVHL tRNA transcripts.

### Quantification of readthrough levels for different 3′ contexts *IN VITRO*

To investigate this amazing phenomenon, we constructed several mRNA fragments containing two stop codons separated by different hexanucleotide sequences (Figure 1C). We chose most frequent stop codon, UAA, to be the second stop codon in order to exclude secondary readthrough and to estimate the level of first stop codon readthrough. The 3′ context of the second stop codon was A-rich AAG CUU; this ensured efficient translation termination according to our data (43). As internal contexts, we chose six different 3′ sequences annotated earlier as signals provoking readthrough (Table 1). In addition to the two weak sequences described above, we cloned a third weak context containing a probable weak CU dinucleotide followed by UAA stop codon, to be able to detect a +2 frameshift during the stop codon readthrough. A +1 frameshift during stop codon readthrough could be detected using the weak 1 context that contains UAG stop codon in the +1 position. Additionally, we cloned two other known weak 3′ contexts following the UAG stop codon: TMV from the tobacco mosaic virus (30) and Dyst that corresponds to the sequence followed the nonsense mutation 651d in human mRNA dystrophin transcript variant Dp4271 (53). The Dyst context also contains the internal stop codon UAA that made possible another +2 frameshift.

To estimate stop codon readthrough efficiency, we performed toe-printing assays of the preTCs assembled at corresponding mRNAs. We significantly increased the total amount of tRNA in the reaction, to exclude +3 nucleotide pause from appearing after the readthrough. In all tested weak 3′ stop codon contexts, part of the ribosomes passed through the first stop codon and paused at the second stop codon, giving the appearance of an additional +9 nucleotide peak (Figure 1D). The ratio of peaks, at the +9 nucleotide and the preTC positions, was used as a measure of the first stop codon readthrough (Figure 1E). The strongest readthrough (∼80%) was observed for the UGA-Weak 1 pair. Two other CU contexts (Weak 2 and Weak 3) also induced pronounced UGA readthrough (∼60-70%). Readthrough of UAG followed by Dyst and TMV contexts were significantly lower (∼20%). It should be noted that we did not observe any readthrough for the standard context (Figure 1D). Additionally, we did not observe any frameshifts during stop codon readthrough (Figure 1D).

### 3′ stop codon contexts do not affect translation termination efficiency

To determine how the chosen weak 3′ contexts affected translation termination, we added eRF1(AGQ) mutant and two variants of eRF3 (eRF3a and eRF3c) to preTCs assembled on various mRNAs containing two stop codons (Figure 1C). eRF1(AGQ) mutant effectively recognises stop codons but does not induce peptide release (54) to exclude additional readthrough during translation termination. Both variants of eRF3 activate eRF1 during translation termination, but differ by the absence of the N-terminal region in eRF3c, which is responsible for binding with poly(A) binding protein (PABP) (45). During stop codon recognition by eRF1+/−eRF3, the ribosome protects additional nucleotides on the mRNA, which can be detected in toe-printing assays as a one or two-nucleotide forward shift of the ribosomal complex (3, 10, 11). The purified preTCs contained mixture of the ribosomal complexes paused at the first (UAG or UGA) and the second (UAA) stop codons. The addition of release factors to this mixture lead to the appearance of peaks corresponding to the termination complex (TC) (Figure 3A).

**Figure 3.**
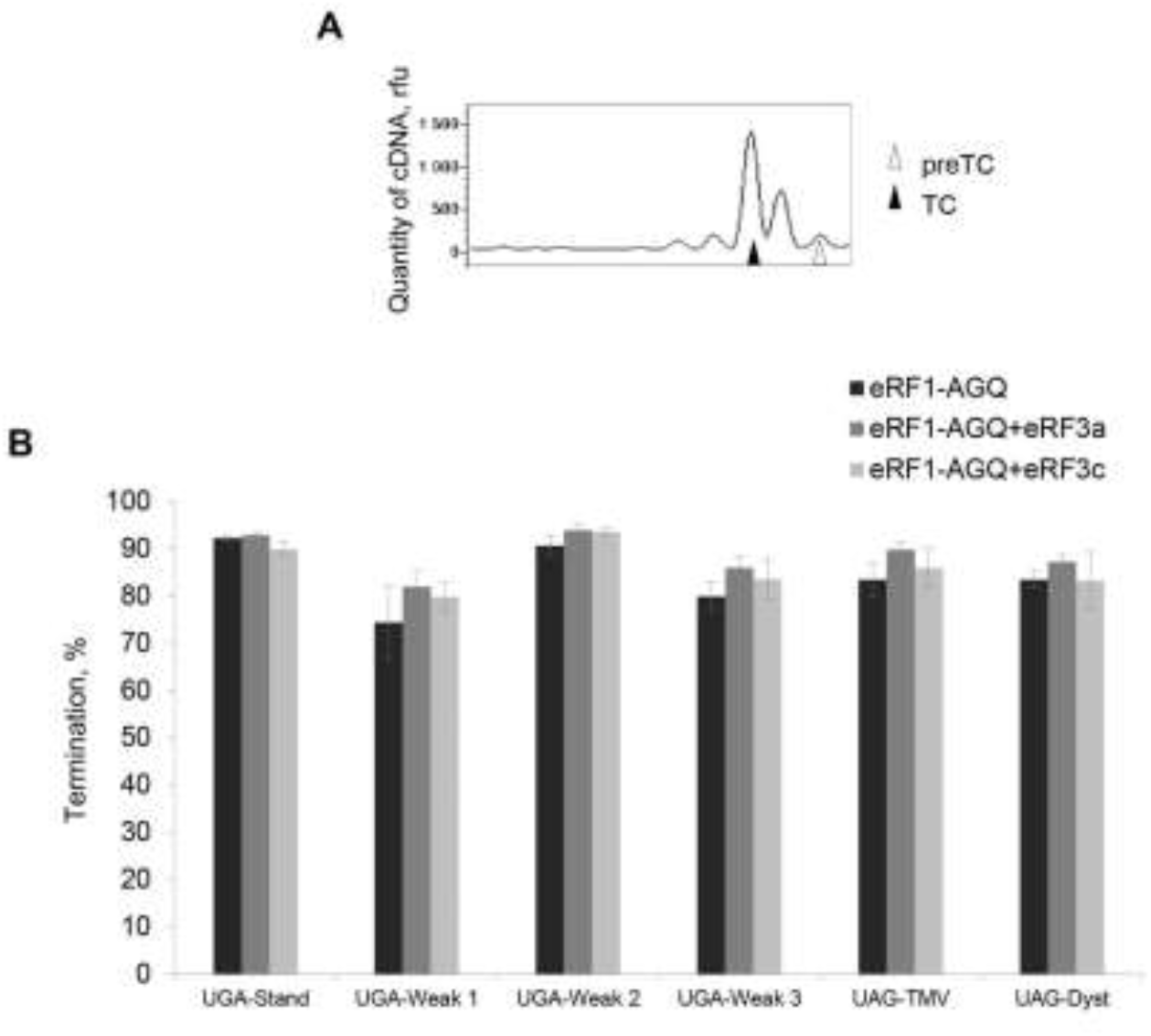
Translation termination in the presence of different stop codon contexts. (A) Example of toe-print analysis of ribosomal complexes assembled with the dual-stop mRNAs containing standard stop codon context and the addition of eRF1(AGQ) and eRF3a. (B) Quantification of the percentage of translation termination at the first stop codon of the dual-stop mRNAs after the addition of eRF1(AGQ) and eRF3a/c, n=3.

We estimated the amount of TCs in different contexts during translation termination (Figure 3B). The percentage of TCs formed at the 1^st^ stop codon at the standard context reached ∼90%. All weak 3′ contexts demonstrated almost equal percentage of TCs (∼ 80%). Therefore, this study revealed that the addition of eRFs ensured high efficiency translation termination, independent of 3′ mRNA sequence.

Our data also shown, that the readthrough occurred without any frameshifting, as we observed by toe-printing only two-nucleotide forward shifts of the ribosomes during termination of translation.

## DISCUSSION

We conducted a study of the impact of stop codon 3′ region on the efficiency of translation termination. Previous studies have shown evidence of programmed stop codon readthrough, which has biological significance (27). We performed an investigation of the mechanism of stop codon readthrough in the presence of the weak 3′ contexts. We, therefore, checked the effect of five reconstituted stop codon 3′ contexts described as weak, on translation termination (Table 1). Surprisingly, the results revealed that the weak contexts do not significantly suppress translation termination, while they induce stop codon readthrough entirely at the elongation stage. We propose that the percentage readthrough is determined by the interaction of the mRNA with the ribosome and that eRFs are not involved. This data is in agreement with previous studies that discuss the mechanism whereby 3′ context impacts readthrough. The earliest approach proposes destabilization of the ribosomal secondary structure owing to the interaction between the rRNA and the mRNA, which leads to an increased probability of the stop codon binding with tRNA than with eRF1 (36). Namy et al. revealed two regions of *S. cerevisiae* rRNA, potentially capable of pairing with 3′ context on mRNA. The first region was found in helix 17 of 18S rRNA (479-510 nt), which is connected to the A-site of the ribosome, and the second region of 18S rRNA was between 1305 and 1318 nt. The mRNA–rRNA base pairing remains to be experimentally tested, however. Recent computational analyses performed by Panek et al. (55) identified mRNA 3′ UTR sequences from 14 eukaryotic species and revealed 18S rRNA complementarity with the first 50 nucleotides of 3′ UTRs, which forms an evolutionary conserved pattern localized around the ribosomal mRNA entry channel. Panek et al. assumed that 18S-3′ UTR base-pairing might act as a mechanism, stabilizing post-termination ribosomal complexes onto mRNA fragment to aid further recycling or to impede their migration. Another recent study, conducted in mammalian cells, also proposed that translation termination efficiency is influenced by interactions of the ribosome with six mRNA nucleotides downstream of the stop codon, occupying the entry channel (40). Interestingly, in that work Cridge et al. showed that among weak contexts the UGA CUA sequence was not responsive to the rise of eRF1 cellular levels, though reducing eRF1 levels by siRNAs dramatically increased the readthrough levels. This agrees with our data, which show that weak contexts mostly influence the readthrough in the absence of eRFs, rather than during the termination stage.

Our results show that the effect of different 3′ contexts on readthrough levels is provoked not by the strength of translation termination, but rather by the ribosome’s capacity to recognize these sequences (Figure 4). It turns out that the ability of eRFs to recognize stop codons and cause peptide release is not affected by the mRNA context. Thus, the efficiency of translation termination is based on an equilibrium between the proportions of ribosomes bound to eRFs and near-cognate tRNAs. This equilibrium shifts back and forth depending on the context of the stop codons.

**Figure 4.**
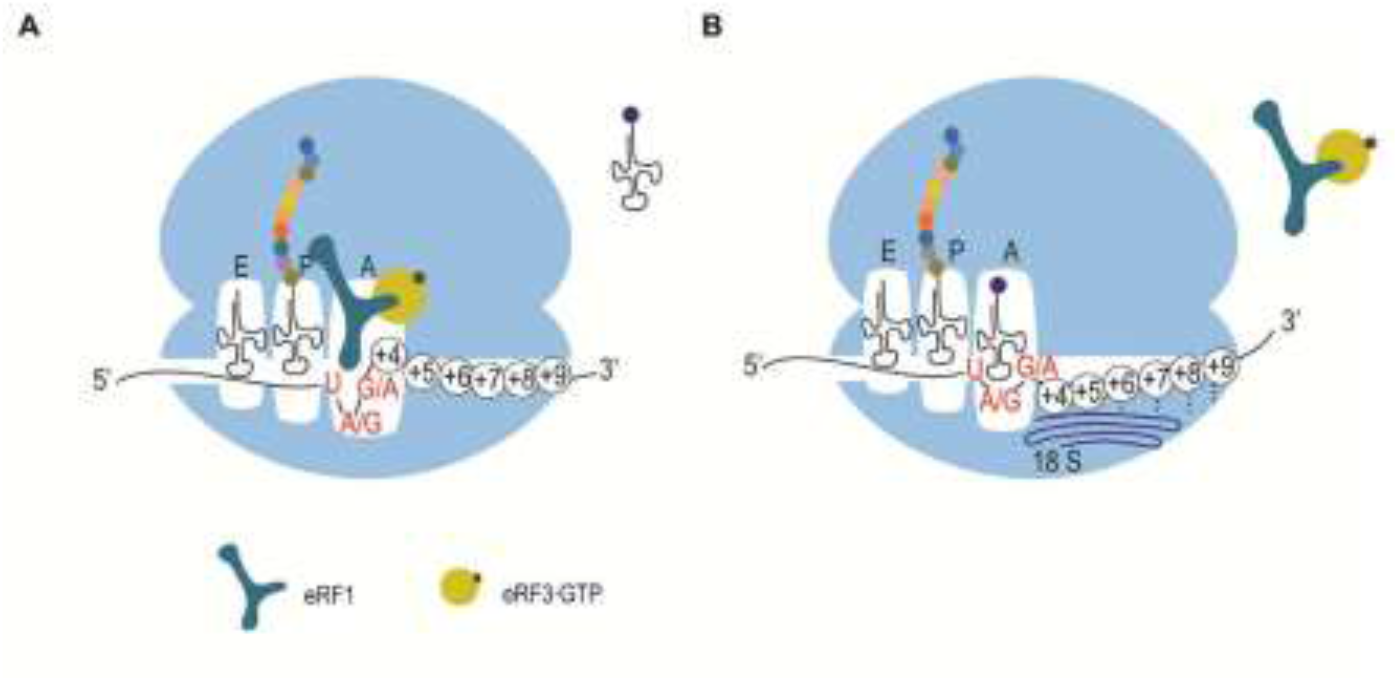
Model of the effect of the 3′ stop codon contexts on translation termination. (A) During translation termination, mRNA with the stop codon followed by random nucleotides undergoes a U-turn-like conformation in the A site, which is preferable for stop codon recognition with eRFs. Near-cognate tRNAs are unable to interact with this structure. (B) Six nucleotides in the weak 3′ stop codon context interact with the rRNA and violate the U-turn like structure of the mRNA in the A-site of the ribosome. This interaction makes the incorporation of near-cognate tRNA preferable to stop codon recognition by eRFs.

Readthrough levels are higher when analysed in pure translation termination systems than in the cell reporter systems. It can reflect much more sophisticated mechanism of balancing between readthrough and translation termination, orchestrated by nucleotide surrounding of stop codon, than could be surmised on the first glance. Obviously, the mechanism of stop codon readthrough in living cells is more difficult and it is regulated by additional factors.

Based on our findings, we propose a model of 3′ context influence on competition between eRF1 and near-cognate tRNA for stop codon recognition (Figure 4). It was shown by groups of Beckman and Ramakrishnan that the +4 nucleotide binds with eRF1 and changes the conformation of the stop codon to form a U-turn like structure of mRNA in the A-site of the ribosome, which is obviously essential for stop codon recognition by eRF1 (10, 11). We assume that this U-turn conformation forms in almost any nucleotide context downstream of the stop codon, which permits the +4 nucleotide to stack with G626 of 18S rRNA (Figure 4A). It is likely that a limited number of specific weak sequences in the 3′ region of the stop codon bonds between base-pairs or stacks with some more extensive regions of 18S rRNA, as was supposed previously (36, 40). We suggest that such interactions may slightly pull +4 nucleotide of mRNA out of the decoding centre, which, then, do not allow the +4 nucleotide to enter the A site and makes mRNA conformation preferable for stop codon recognition by near-cognate tRNA rather than eRF1 (Fig. 4B).

In conclusion, it should be noted that this proposed mechanism allows eRF1 to terminate translation at the overwhelming majority of 3′ stop codon contexts and efficiently enter other stages of translation. Such an approach is reasonable because otherwise any random mutation in this region would lead to the immediate cessation of protein synthesis and a wasting of cell resources. However, the signal impairing translation termination in some cases is needed. This function is realized by a small number of specific sequences capable of interacting with the ribosome.

## ACKNOWLEDGEMENT

We are grateful to Ludmila Frolova for providing us with plasmids encoding release factors, to Tatyana Pestova and Christopher Hellen who provided us with plasmids encoding initiation factors, to Christiane Schaffitzel who provided us with plasmids encoding eRF3a. cDNA fragment analyses were performed by the centre of the collective use “Genome” of EIMB RAS.

## SUPPORTING INFORMATION

**Table S1.**
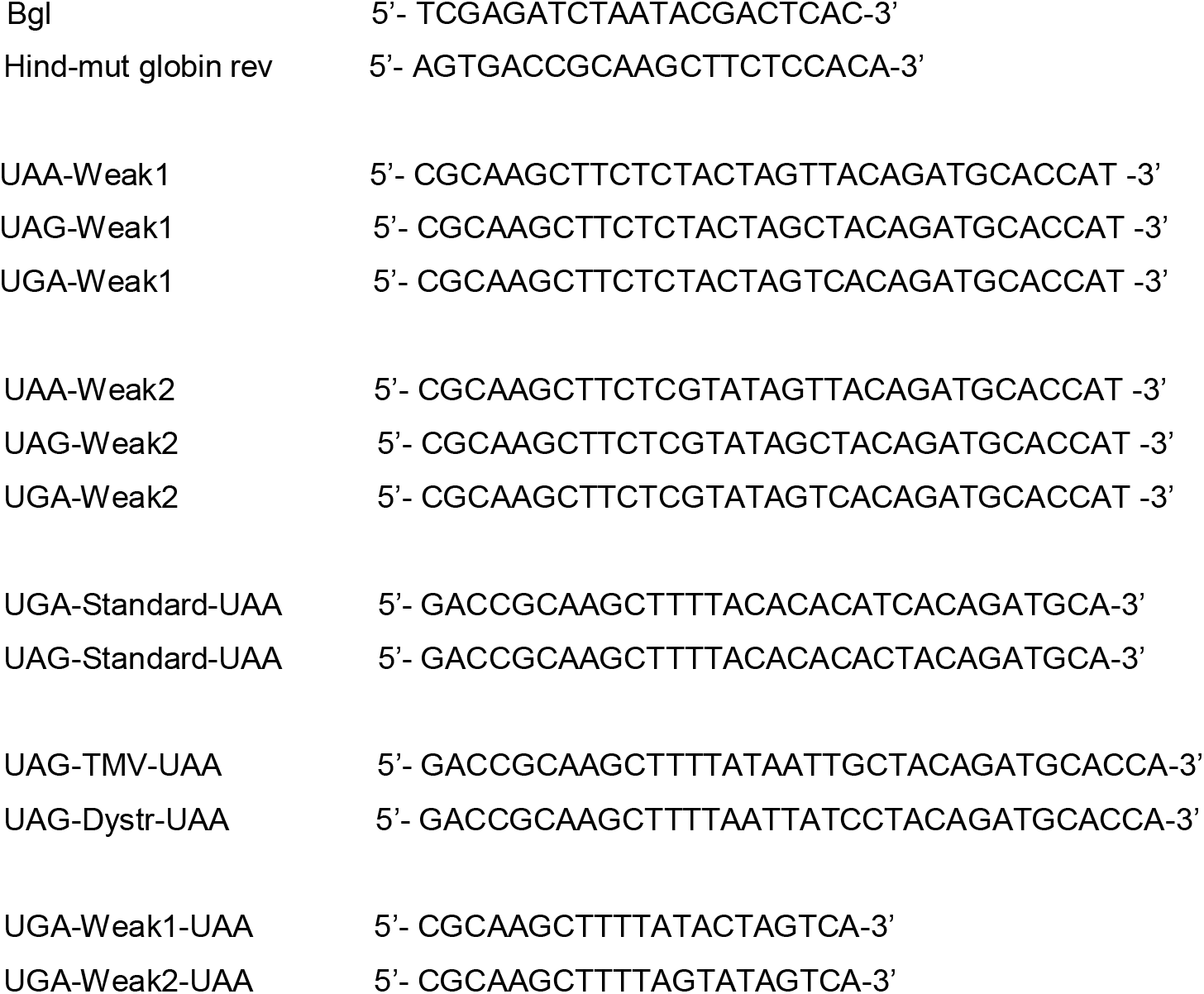
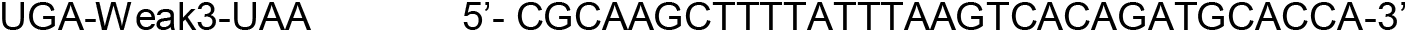
Primers for obtaining MVHL-stop plasmids with various 3’ contexts of stop codons:

**Table S2.** List of cloned constructions.

| Construction | Matrix | Primers | Restriction endonucleases |
| --- | --- | --- | --- |
| pET28a MVHL-UAA- HindIII | pET28a MVHL- UAA | BglII, Hind-mut globin rev | BglII, HindIII |
| UAA Weak1 | pET28a MVHL- UAA-HindIII | UAA-Weak1, Bgl | BglII, HindIII |
| UAG Weak1 | pET28a MVHL- UAA-HindIII | UAG-Weak1, Bgl | BglII, HindIII |
| UGA Weak1 | pET28a MVHL- UAA-HindIII | UGA-Weak1, Bgl | BglII, HindIII |
| UAA Weak2 | pET28a MVHL- UAA-HindIII | UAA-Weak2, Bgl | BglII, HindIII |
| UAG Weak2 | pET28a MVHL- UAA-HindIII | UAG-Weak2, Bgl | BglII, HindIII |
| UGA Weak2 | pET28a MVHL- UAA-HindIII | UGA-Weak2, Bgl | BglII, HindIII |
| UAG Standard-UAA | pET28a MVHL- UAA-HindIII | UAG-Standard- UAA, Bgl | BglII, HindIII |
| UGA Standard-UAA | pET28a MVHL- UAA-HindIII | UGA-Standard- UAA, Bgl | BglII, HindIII |
| UAG TMV-UAA | UAG Standard-UAA | UAG-TMV-UAA, Bgl | BglII, HindIII |
| UAG Dystro-UAA | UAG Standard-UAA | UAG-Dystro-UAA, Bgl | BglII, HindIII |
| UGA Weak1-UAA | UGA Weak1 context | UGA-Weak1-UAA, Bgl | BglII, HindIII |
| UGA Weak2-UAA | UGA Weak2 context | UGA-Weak2-UAA, Bgl | BglII, HindIII |
| UGA Weak3-UAA | UGA Weak1 context | UGA-Weak3-UAA, Bgl | BglII, HindIII |

## REFERENCES

1. Zhouravleva, G., Frolova, L., Le Goff, X., Le Guellec, R., Inge-Vechtomov, S., Kisselev, L., and Philippe, M. (1995) Termination of translation in eukaryotes is governed by two interacting polypeptide chain release factors, eRF1 and eRF3. The EMBO journal. 14, 4065–72

2. Stansfield, I., Jones, K. M., Ter-Avanesyan, M. D., and Tuite5, M. F. (1995) The products of the SUP45 (eRF1) and SUP35 genes interact to mediate translation termination in Saccharomyces cerevisiae. The EMBO Journal. 14, 4365–4373

3. Alkalaeva, E. Z., Pisarev, A. V., Frolova, L. Y., Kisselev, L. L., and Pestova, T. V. (2006) In Vitro Reconstitution of Eukaryotic Translation Reveals Cooperativity between Release Factors eRF1 and eRF3. Cell. 125, 1125–1136

4. Nakamura, Y., Ito, K., and Ehrenberg, M. (2000) Mimicry grasps reality in translation termination. Cell. 101, 349–352

5. Voorhees, R. M., Schmeing, T. M., Kelley, A. C., and Ramakrishnan, V. (2010) The mechanism for activation of GTP hydrolysis on the ribosome. Science (New York, N.Y.). 330, 835–838

6. Des Georges, A., Hashem, Y., Unbehaun, A., Grassucci, R. A., Taylor, D., Hellen, C. U. T., Pestova, T. V., and Frank, J. (2014) Structure of the mammalian ribosomal pre-termination complex associated with eRF1???eRF3???GDPNP. Nucleic Acids Research. 42, 3409–3418

7. Bertram, G., Bell, H. A., Ritchie, D. W., Fullerton, G., and Stansfield, I. (2000) Terminating eukaryote translation: Domain 1 of release factor eRF1 functions in stop codon recognition. RNA. 6, 1236–1247

8. Ito, K., Frolova, L., Seit-Nebi, A., Karamyshev, A., Kisselev, L., and Nakamura, Y. (2002) Omnipotent decoding potential resides in eukaryotic translation termination factor eRF1 of variant-code organisms and is modulated by the interactions of amino acid sequences within domain 1. Proceedings of the National Academy of Sciences of the United States of America. 99, 8494–9

9. Feng, T., Yamamoto, A., Wilkins, S. E., Sokolova, E., Yates, L. A., Münzel, M., Singh, P., Hopkinson, R. J., Fischer, R., Cockman, M. E., Shelley, J., Trudgian, D. C., Schödel, J., McCullagh, J. S. O., Ge, W., Kessler, B. M., Gilbert, R. J., Frolova, L. Y., Alkalaeva, E., Ratcliffe, P. J., Schofield, C. J., and Coleman, M. L. (2014) Optimal Translational Termination Requires C4 Lysyl Hydroxylation of eRF1. Molecular Cell. 10.1016/j.molcel.2013.12.028

10. Brown, A., Shao, S., Murray, J., Hegde, R. S., and Ramakrishnan, V. (2015) Structural basis for stop codon recognition in eukaryotes. Nature. 10.1038/nature14896

11. Matheisl, S., Berninghausen, O., Becker, T., and Beckmann, R. (2015) Structure of a human translation termination complex. Nucleic Acids Research. 10.1093/nar/gkv909

12. Shao, S., Murray, J., Brown, A., Taunton, J., Ramakrishnan, V., and Hegde, R. S. (2016) Decoding Mammalian Ribosome-mRNA States by Translational GTPase Complexes. Cell. 10.1016/j.cell.2016.10.046

13. McCaughan, K. K., Brown, C. M., Dalphin, M. E., Berry, M. J., and Tate, W. P. (1995) Translational termination efficiency in mammals is influenced by the base following the stop codon. Proceedings of the National Academy of Sciences of the United States of America. 92, 5431–5435

14. Keeling, K. M., Xue, X., Gunn, G., and Bedwell, D. M. (2014) Therapeutics Based on Stop Codon Readthrough. Annual Review of Genomics and Human Genetics. 15, 371–394

15. Loughran, G., Chou, M. Y., Ivanov, I. P., Jungreis, I., Kellis, M., Kiran, A. M., Baranov, P. V., and Atkins, J. F. (2014) Evidence of efficient stop codon readthrough in four mammalian genes. Nucleic Acids Research. 42, 8928–8938

16. Schueren, F., Lingner, T., George, R., Hofhuis, J., Dickel, C., Gärtner, J., and Thoms, S. (2014) Peroxisomal lactate dehydrogenase is generated by translational readthrough in mammals. eLife. 3, e03640

17. Jungreis, I., Lin, M. F., Spokony, R., Chan, C. S., Negre, N., Victorsen, A., White, K. P., and Kellis, M. (2011) Evidence of abundant stop codon readthrough in Drosophila and other metazoa. Genome Research. 21, 2096–2113

18. Dunn, J. G., Foo, C. K., Belletier, N. G., Gavis, E. R., and Weissman, J. S. (2013) Ribosome profiling reveals pervasive and regulated stop codon readthrough in Drosophila melanogaster. eLife. 10.7554/eLife.01179

19. Freitag, J., Ast, J., and Bölker, M. (2012) Cryptic peroxisomal targeting via alternative splicing and stop codon read-through in fungi. Nature. 485, 522–525

20. Chittum, H. S., Lane, W. S., Carlson, B. A., Roller, P. P., Lung, F. D. T., Lee, B. J., and Hatfield, D. L. (1998) Rabbit β-globin is extended beyond its UGA stop codon by multiple suppressions and translational reading gaps. Biochemistry. 37, 10866–10870

21. Yamaguchi, Y., Hayashi, A., Campagnoni, C. W., Kimura, A., Inuzuka, T., and Baba, H. (2012) L-MPZ, a novel isoform of myelin P0, is produced by stop codon readthrough. Journal of Biological Chemistry. 287, 17765–17776

22. Dunn, J. G., Foo, C. K., Belletier, N. G., Gavis, E. R., and Weissman, J. S. (2013) Ribosome profiling reveals pervasive and regulated stop codon readthrough in Drosophila melanogaster. eLife. 2013, 1–32

23. Eswarappa, S. M., Potdar, A. A., Koch, W. J., Fan, Y., Vasu, K., Lindner, D., Willard, B., Graham, L. M., Dicorleto, P. E., and Fox, P. L. (2014) Programmed translational readthrough generates antiangiogenic VEGF-Ax. Cell. 157, 1605–1618

24. Loughran, G., Jungreis, I., Tzani, I., Power, M., Dmitriev, R. I., Ivanov, I. P., Kellis, M., and Atkins, J. F. (2018) Stop codon readthrough generates a C-terminally extended variant of the human vitamin D receptor with reduced calcitriol response. The Journal of biological chemistry. 10.1074/jbc.M117.818526

25. Brar, G. A. (2016) Beyond the Triplet Code: Context Cues Transform Translation. Cell. 167, 1681–1692

26. Dabrowski, M., Bukowy-Bieryllo, Z., and Zietkiewicz, E. (2015) Translational readthrough potential of natural termination codons in eucaryotes – The impact of RNA sequence. RNA Biology. 12, 950–958

27. Baranov, P. V, Atkins, J. F., and Yordanova, M. M. (2015) Augmented genetic decoding: global, local and temporal alterations of decoding processes and codon meaning. Nature reviews. Genetics. 16, 517–29

28. Bertram, G., Innes, S., Minella, O., Richardson, J., and Stansfield, I. (2001) Endless possibilities: translation termination and stop codon recognition. Microbiology (Reading, England). 147, 255–69

29. Pedersen, W. T., and Curran, J. F. (1991) Effects of the nucleotide 3??? to an amber codon on ribosomal selection rates of suppressor tRNA and release factor-1. Journal of Molecular Biology. 219, 231–241

30. Skuzeski, J. M., Nichols, L. M., Gesteland, R. F., and Atkins, J. F. (1991) The signal for a leaky UAG stop codon in several plant viruses includes the two downstream codons. Journal of Molecular Biology. 218, 365–373

31. Li, G., and Rice, C. M. (1993) The signal for translational readthrough of a UGA codon in Sindbis virus RNA involves a single cytidine residue immediately downstream of the termination codon. Journal of virology. 67, 5062–5067

32. Bonetti, B., Fu, L., Moon, J., and Bedwell, D. M. (1995) The efficiency of translation termination is determined by a synergistic interplay between upstream and downstream sequences in Saccharomyces cerevisiae. Journal of molecular biology. 251, 334–45

33. Brown, C. M., Stockwell, P. A., Trotman, C. N., and Tate, W. P. (1990) Sequence analysis suggests that tetra-nucleotides signal the termination of protein synthesis in eukaryotes. Nucleic acids research. 18, 6339–45

34. Tate, W. P., and Mannering, S. A. (1996) MicroReview Three, four or more?: the translational stop signal at length. Molecular microbiology. 21, 213–219

35. Poole, E. S., Major, L. L., Mannering, S. A., and Tate, W. P. (1998) Translational termination in Escherichia coli: Three bases following the stop codon crosslink to release factor 2 and affect the decoding efficiency of UGA-containing signals. Nucleic Acids Research. 26, 954–960

36. Namy, O., Hatin, I., and Rousset, J. P. (2001) Impact of the six nucleotides downstream of the stop codon on translation termination. EMBO Reports. 2, 787–793

37. Cavener, D. R., and Ray, S. C. (1991) Eukaryotic start and stop translation sites. Nucleic acids research. 19, 3185–92

38. Cridge, A. G., Major, L. L., Mahagaonkar, A. A., Poole, E. S., Isaksson, L. A., and Tate, W. P. (2006) Comparison of characteristics and function of translation termination signals between and within prokaryotic and eukaryotic organisms. Nucleic acids research. 34, 1959–73

39. Williams, I., Richardson, J., Starkey, A., and Stansfield, I. (2004) Genome-wide prediction of stop codon readthrough during translation in the yeast Saccharomyces cerevisiae. Nucleic Acids Research. 32, 6605–6616

40. Cridge, A. G., Crowe-Mcauliffe, C., Mathew, S. F., and Tate, W. P. (2018) Eukaryotic translational termination efficiency is influenced by the 3 nucleotides within the ribosomal mRNA channel. Nucleic Acids Research

41. Beznosková, P., Wagner, S., Jansen, M. E., Von Der Haar, T., and Valášek, L. S. (2015) Translation initiation factor eIF3 promotes programmed stop codon readthrough. Nucleic Acids Research. 43, 5099–5111

42. Loenarz, C., Sekirnik, R., Thalhammer, A., Ge, W., Spivakovsky, E., Mackeen, M. M., McDonough, M. A., Cockman, M. E., Kessler, B. M., Ratcliffe, P. J., Wolf, A., and Schofield, C. J. (2014) Hydroxylation of the eukaryotic ribosomal decoding center affects translational accuracy. Proceedings of the National Academy of Sciences. 111, 4019–4024

43. Sokolova, E. E., Vlasov, P. K., Egorova, T. V., Shuvalov, A. V., and Alkalaeva, E. Z. (2020) The Influence of A/G Composition of 3’ Stop Codon Contexts on Translation Termination Efficiency in Eukaryotes. Molekuliarnaia biologiia. 10.31857/S0026898420050080

44. Alkalaeva, E., Eliseev, B., Ambrogelly, A., Vlasov, P., Kondrashov, F. A., Gundllapalli, S., Frolova, L., Söll, D., and Kisselev, L. (2009) Translation termination in pyrrolysine-utilizing archaea. FEBS Letters. 583, 3455–3460

45. Ivanov, A., Mikhailova, T., Eliseev, B., Yeramala, L., Sokolova, E., Susorov, D., Shuvalov, A., Schaffitzel, C., and Alkalaeva, E. (2016) PABP enhances release factor recruitment and stop codon recognition during translation termination. Nucleic Acids Research. 10.1093/nar/gkw635

46. Kryuchkova, P., Grishin, A., Eliseev, B., Karyagina, A., Frolova, L., and Alkalaeva, E. (2013) Two-step model of stop codon recognition by eukaryotic release factor eRF1. Nucleic Acids Research. 41, 4573–4586

47. Susorov, D., Mikhailova, T., Ivanov, A., Sokolova, E., and Alkalaeva, E. (2015) Stabilization of eukaryotic ribosomal termination complexes by deacylated tRNA. Nucleic Acids Research. 43, 3332–3343

48. Egorova, T., Sokolova, E., Shuvalova, E., Matrosova, V., Shuvalov, A., and Alkalaeva, E. (2019) Fluorescent toeprinting to study the dynamics of ribosomal complexes. Methods. 10.1016/j.ymeth.2019.06.010

49. Susorov, D., Mikhailova, T., Ivanov, A., Sokolova, E., and Alkalaeva, E. (2015) Stabilization of eukaryotic ribosomal termination complexes by deacylated tRNA. Nucleic Acids Research. 10.1093/nar/gkv171

50. Ivanov, A., Mikhailova, T., Eliseev, B., Yeramala, L., Sokolova, E., Susorov, D., Shuvalov, A., Schaffitzel, C., and Alkalaeva, E. (2016) PABP enhances release factor recruitment and stop codon recognition during translation termination. Nucleic Acids Research. 44, 7766–7776

51. Blanchet, S., Cornu, D., Argentini, M., and Namy, O. (2014) New insights into the incorporation of natural suppressor tRNAs at stop codons in Saccharomyces cerevisiae. Nucleic Acids Research. 42, 10061–10072

52. Beznosková, P., Ová, S. G. Š., Shivaya, L. E. O. Š., and Ek, V. Š. (2016) Rules of UGA-N decoding by near-cognate tRNAs and analysis of readthrough on short uORFs in yeast. 10.1261/rna.054452.115.

53. Bidou, L., Hatin, I., Perez, N., Allamand, V., Panthier, J., and Rousset, J. (2004) Premature stop codons involved in muscular dystrophies show a broad spectrum of readthrough efficiencies in response to gentamicin treatment. Gene Therapy. 11, 619–627

54. Seit-Nebi, A., Frolova, L., Justesen, J., and Kisselev, L. (2001) Class-1 translation termination factors: Invariant GGQ minidomain is essential for release activity and ribosome binding but not for stop codon recognition. Nucleic Acids Research. 29, 3982–3987

55. Pánek, J., Kolář, M., Herrmannová, A., and Valášek, L. S. (2016) A systematic computational analysis of the rRNA–3′ UTR sequence complementarity suggests a regulatory mechanism influencing post-termination events in metazoan translation. RNA. 22, 957–967

